# Targeting HIV-1 with CRISPR/Cas9 delivered by retargeted adenoviruses effectively suppresses viral replication

**DOI:** 10.1101/2023.12.18.572146

**Authors:** Sarah Klinnert, Patrick C. Freitag, Andreas Plückthun, Karin J. Metzner

**Author notes:** Corresponding author: Karin J. Metzner, M.D. University Hospital Zurich Department of Infectious Diseases and Hospital Epidemiology Rämistrasse 100 CH-8091 Zurich.

## Abstract

Integrated, intact, latent HIV-1 viruses in infected cells are the main obstacle to curing HIV-1 infections. Targeted inactivation of HIV-1 proviruses with CRISPR/Cas9 is a promising strategy to eradicate HIV-1. In addition, CRISPR/Cas9 is able to target replicating HIV-1 and could be used as a therapy during productive infection.

Here, we combine the CRISPR/Cas9 system with a novel adenovirus (Ad) targeted delivery technology to test it as a therapeutic approach to inhibit HIV-1. First, we selected six HIV-1-specific gRNAs targeting the HIV-1 LTRs and the *gag* gene and tested their efficacy in inhibiting HIV-1 virion production in an HEK 293T cell co-transfection screen. The gRNA-TAR showed the most robust and potent inhibition of HIV-1 by >99% alone or in combination with the gRNA-p24, which induced a ∼1 kb deletion between both gRNA target sites in HIV-1 DNA. Delivery of this dual gRNA-TAR/p24 CRISPR/Cas9 system with CD3-CD28-IL2-retargeted Ads was highly effective, transducing 62.3±23.3% of cells and suppressing HIV-1 replication by 88.0±4.5% in primary CD4^+^ T cells from three independent donors.

Our dual gRNA-TAR/p24-CRISPR/Cas9-Ad strategy represents a novel therapeutic approach to effectively inhibit HIV-1 in a highly HIV-1 and T cell-specific manner.

## Introduction

Integration of its viral genome into the host cell genome is a key characteristic of the virus family *retroviridae* to which the human immunodeficiency virus type 1 (HIV-1) belongs. This integrated HIV-1 genome, also called provirus, serves as a template for the transcription of viral RNAs and hence is vital for virus replication. In a small subset of HIV-1 infected CD4^+^ cells intact provirus becomes latent, meaning it is transcriptionally silent, but it can be reactivated and reestablish active virus replication. Due to these HIV-1 latently infected cells also known as the latent reservoir, HIV-1 cannot be cured and combination antiretroviral therapy (cART), which effectively inhibits virus replication but does not affect the latent reservoir, has to be taken life-long ^1-3^.

Genome editing technologies have been investigated regarding their applicability to specifically inactivate HIV-1 provirus. Designable DNA-binding nucleases such as zinc finger nucleases (ZFNs) and transcription activator-like effector nucleases (TALENs) exhibited effective excision and elimination of HIV-1 provirus from the host cell genome ^4-7^. Both technologies show high editing efficiencies and a seemingly unlimited targeting range; additionally TALENs in particular show low cytotoxicity and off-target effects. The newest genome editing technology, the CRISPR/Cas system, which consists of an RNA-guided Cas nuclease, additionally provides the advantages of easy, timely and cost-effective application and change or addition of new target sites by guide RNA (gRNA) design ^8^.

HIV-1 targeting with CRISPR/Cas is one of the major HIV-1 eradication strategies and has been investigated in numerous studies using *Staphylococcus pyogenes* Cas9, *Staphylococcus aureus* Cas9, Cas12a or Cas13a, and targeting different regulatory or structural genome region such as the LTR, *gag*, *pol*, *tat* and *rev* genes ^9^. Furthermore, the CRISPR/Cas system can also be harnessed as a therapy during acute HIV-1 infection to inhibit and suppress virus replication while simultaneously targeting integrated HIV-1 proviruses ^10,11^. Priorities for the development of an effective and safe CRISPR/Cas therapy against HIV-1 are the identification of efficacious gRNA target sites that are highly conserved among divergent viral strains, multiplexing of gRNAs and an effective delivery method.

In this study, we combined the CRISPR/Cas9 genome editing technology and a novel T cell targeted adenovirus (Ad) vector delivery system to target HIV-1. First, we systematically tested different gRNA target sites in the HIV-1 genome to identify the most effective gRNAs for editing and suppression of HIV-1 expression by the CRISPR/Cas9 system. Second, we used novel CD3-CD28-IL2 retargeting adenovirus vectors to deliver the most potent gRNA pair very efficiently into primary T cells.

## Materials and Methods

### Plasmids

pSpCas9(BB)-2A-GFP (PX458) was a gift from Feng Zhang (Addgene plasmid #48138) ^12^. Selected HIV-1 specific and control gRNAs (Table 1) were cloned into the pSpCas9(BB)-2A-GFP plasmid using the sgRNA cloning protocol from S. Konermann, Zhang lab, 2014 ^13^. The HIV-1 full-length plasmids pYK-JRCSF and pBR-NL4-3 Nef^+^ IRES eGFP were obtained through the NIH HIV Reagent Program, Division of AIDS, NIAID, NIH: Human immunodeficiency virus 1 (HIV-1), strain JR-CSF infectious molecular clone (pYK-JRCSF), was contributed by Irvin SY Chen and Yoshio Koyanagi ^14^. HIV-1_NL4-3 Nef+ IRES eGFP_ (clone 11349) co-expressing Nef and eGFP from a single bicistronic RNA, was contributed by Frank Kirchhoff ^15^. All plasmids were verified by sequencing.

**Table 1.**
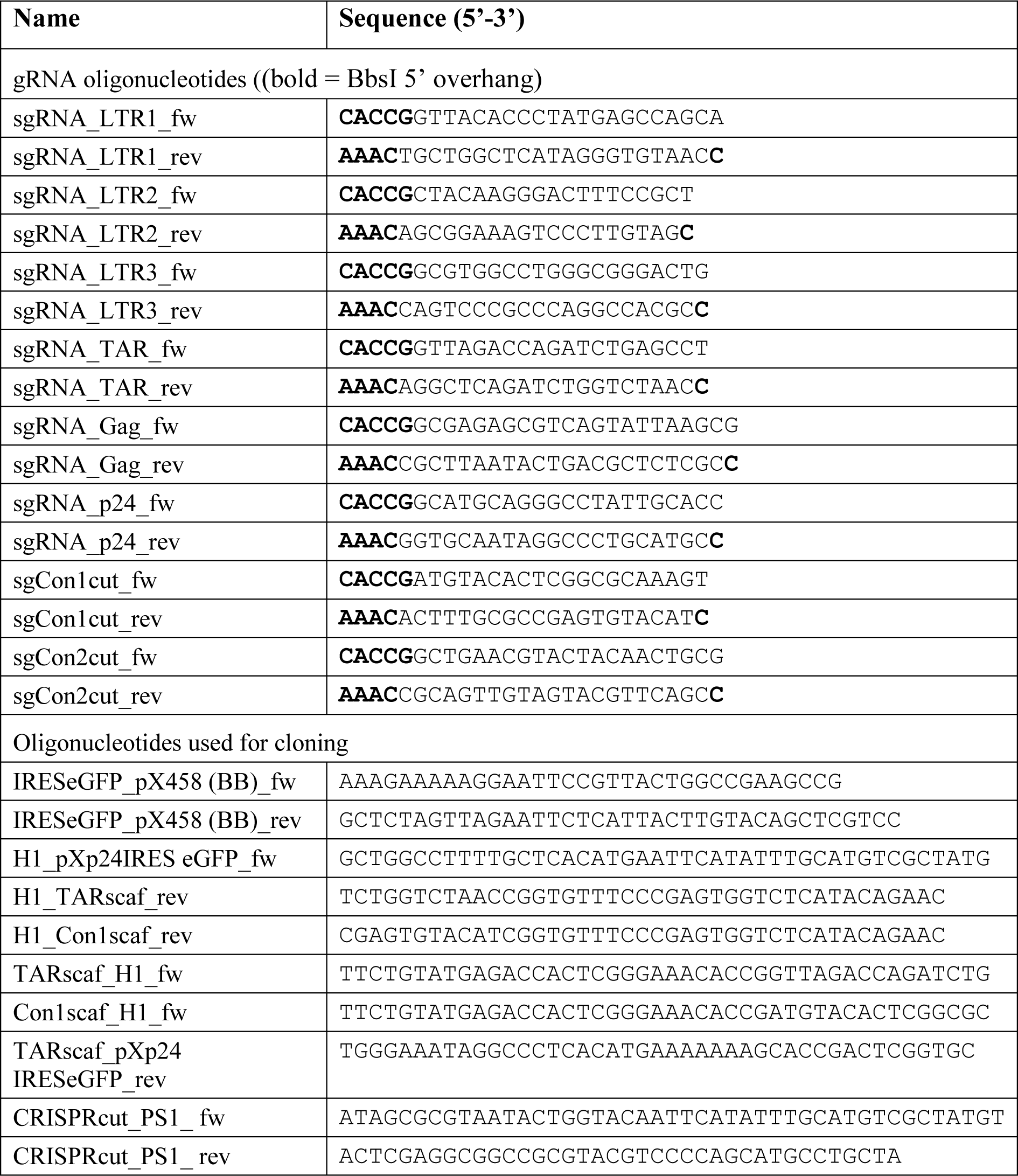
Oligonucleotides.

### Cloning of adenovirus vectors

To delete the T2A-GFP cassette in pSpCas9(BB)-2A-GFP (PX458), the plasmid was cut at its EcoRI restriction sites flanking this T2A-GFP cassette, and 55 ng of the digested plasmid was religated using 1 U of T4 Ligase (Thermo Scientific) resulting in a plasmid termed pSpCas9(BB). The gRNA–p24 and gRNA–Con2 were cloned separately into the pSpCas9(BB) plasmid as mentioned above and the resulting plasmid U6-gRNA-p24-pSpCas9(BB) was cut at the EcoRI restriction site. An IRES-eGFP sequence amplified from a donor plasmid containing the required IRES-eGFP cassette with the primers IRESeGFP_pX458(BB)_fw and IRESeGFP_pX458(BB)_rev containing homologous overlaps with the EcoRI site in the EcoRI cut pSpCas9(BB)-gRNA-p24, was cloned into it using the In-Fusion® HD Cloning Kit (Takara) as described in the manufacturer’s manual. The T2A-GFP cassette was replaced by the IRES-eGFP cassette because the former showed weak expression of GFP and we aimed to monitor Cas9 expression through the GFP reporter.

To add another gRNA expression cassette two sequences were amplified; first, the H1 promoter was amplified from a donor plasmid using the primers H1_pXp24IRESeGFP_fw and H1_TARscaf_rev. Second, the gRNA–TAR and gRNA scaffold sequence was amplified from pSpCas9(BB)-2A-GFP-gRNA-TAR using the primers TARscaf_H1_fw and TARscaf_ pXp24IRESeGFP_rev. Both amplified sequenced contained homologous overlaps with each other and the PciI cut site of the digested backbone U6-gRNA-p24-pSpCas9(BB)-IRES-eGFP. All three DNA fragments were assembled using the In-Fusion® HD Cloning Kit. This resulted in the H1-gRNA-TAR-U6-gRNA-p24-pSpCas9(BB)-IRES-eGFP plasmid. The same was done for the U6-gRNA-Con2-pSpCas9(BB)-IRES-eGFP plasmid using the primers H1_pXp24IRESeGFP_fw and H1_Con1scaf_rev to amplify the H1 promoter, and Con1scaf_H1_fw and TARscaf_ pXp24IRESeGFP_rev to amplify the gRNA-Con1 and gRNA scaffold. This resulted in the H1-gRNA-Con1-U6-gRNA-Con2-pSpCas9(BB)-IRES-eGFP plasmid.

The CRISPR/Cas9-IRES-eGFP expression cassettes containing either the two HIV-1 specific gRNAs, TAR and p24, or control gRNAs, Con1 and Con2, were cloned into the pShuttle (PS-1) vector of the AdEasy adenoviral vector system (Agilent Technologies). The entire cassette was amplified with the primers CRISPRcut_PS1_fw and CRISPRcut_PS1_rev and assembled with the KpnI digested PS-1 plasmid using In-Fusion® HD Cloning. Cloning from the respective PS-1 vectors into pAdEasy-1_HVR7 adenoviral vector was performed according to the manufacturer’s instructions. The pAdEasy-1_HVR7 adenoviral vector is a modified version of the original pAdEasy-1, which reduces liver infection ^16^. HAdV5^HVR7^ recombinant adenoviruses with the respective HIV-1 specific dual gRNA CRISPR/Cas9 expression cassette (Ad-CRISPR-HIV1) or control dual gRNA expression cassette (Ad-CRISPR-Con) were produced by Vector Biolabs (Malvern, PA, USA).

All amplified PCR fragments for cloning were verified on an agarose gel regarding the correct size before extraction and purification for cloning. All intermediate and final plasmids were confirmed by sequencing. All oligonucleotides were purchased from Microsynth, Switzerland, and are listed in table 1.

### Cell culture and transfection

HEK 293T cells ^17^ and TZM-bl cells from J. C. Kappes and X. Wu ^18^were obtained through the NIH AIDS Reagent Programme, Division of AIDS, NIAID, NIH. Cells were maintained in Dulbecco’s Modified Eagle Medium (DMEM) supplemented with 10% heat-inactivated fetal bovine serum (FBS) and 1% penicillin-streptomycin (10,000 units/ml penicillin, 10 mg/ml streptomycin, Gibco) (P/S) at 37 °C and 5% CO_2_. HEK 293T cells were co-transfected with the pSpCas9(BB)-2A-GFP (pX458) containing the Cas9 and different HIV-1 gRNAs, and pYK-JRCSF HIV-1 expression plasmid in a 5:1 mass ratio with Lipofectamine 2000 (Life Technologies) according to the manufacturer’s manual.

### TZM-bl luciferase assay

Analysis of infectious HIV-1 virus particles was achieved by plating cell-free supernatants in a 96-well plate and 2.5×10^4^ TZM-bl cells/well were seeded with it in a total volume of 200 µl medium containing 5 µl Diethylaminoethyl (DEAE)-Dextran. After 24 or 48 h, luciferase expression (relative light units, RLU) was measured with the Bright-Glo™ Luciferase Assay System (Promega).

For analysis of supernatants from HEK 293T cells co-transfected with CRISPR/Cas9, gRNA and HIV-1 JR-CSF expression plasmids 50 µl cell-free supernatant was used; and for Ad-CRISPR transduced and HIV-1_NL4-3-GFP Nef+_ infected primary CD4^+^ T cells 20 µl of each sample were used.

### HIV-1 p24 capsid enzyme-linked immunosorbent assay (ELISA)

For p24 ELISA, 50 µl of cell-free supernatant from cells co-transfected with CRISPR/Cas9, gRNA and HIV-1 JR-CSF expression plasmids was inactivated with 1% Empigen and p24 antigen was quantified using an in-house p24 ELISA assay ^19^.

### InDel PCR and InDel NGS analysis

HEK 293T cells co-transfected with CRISPR/Cas9, gRNA and HIV-1 JR-CSF expression plasmids as described above were centrifuged for 5 minutes (min) at 600 × g 48 h post-transfection. Subsequently, DNAse treatment with 30 U RNase-free DNase I (Roche) per sample in PBS was performed for 20 min at room temperature before inactivating the enzyme at 70°C for 5 min. Total genomic DNA was isolated from DNase-treated cells using the DNeasy® Blood & Tissue Kit (Qiagen) according to the manufacturer’s manual. A PCR was performed using primers that flank the gRNA–TAR and –p24 target sites (5LTR_seq_TZMbl_fw1 (P1) = 5’-GGAAGGGCTAATTCACTCCC-3’; gag 1765 rc (P2) = 5’-CATTTTGGACCAACAAGGTTTCTGTC-3’). The PCR reaction contained 1.25 U JumpStart^TM^ Taq DNA Polymerase (Sigma-Aldrich), 1× PCR Buffer (Sigma-Aldrich), 3 mM MgCl_2_, 1 mM dNTPs, 0.2 µM of each primer and 200 ng of isolated DNA. Cycling conditions were 2 min at 94°C for initial denaturation, followed by 40 cycles of 30 seconds (sec) at 94°C, 30 sec at 56°C and 2 min at 72°C, and a final extension of 5 min at 72°C. After the PCR run 2 µl of the PCR reaction was run on a 1% agarose gel containing ethidium bromide and the GeneRuler 1 kb Plus DNA ladder (Thermo Scientific) was used as a DNA size reference. The remaining PCR reaction of samples treated with a single gRNA was subjected to purification with AMPure XP reagent (Beckman Coulter) according to the manufacturer’s manual using AMPure beads. The remaining PCR reaction of the sample with gRNA–TAR and –p24 combined was run on a 1% agarose gel containing ethidium bromide and the two expected bands were cut out separately and purified using the Nucleospin^®^ Gel and PCR Clean-up kit (Macherey-Nagel) according to the manufacturer’s manual. Afterwards, purified PCR products were subjected to next-generation sequencing (NGS). NGS results were analyzed for InDels using a Python script, which performed the following analysis in chronological order: minimap2 (version 2.17-r941), map reads to given reference, samtools (version 1.7), extract mapped reads and sort them, lofreq (version 2.1.5), call variants including indels.

### Next generation sequencing and analysis

Amplicon preparation for next-generation sequencing using Illumina Nextera XT (Illumina) for 1×150 base pairs (bp) was performed according to the manufacturer’s manual followed by sequencing on a MiSeq system. NGS results were analyzed using the CLC Genomics Workbench Version 22 (Qiagen Digital Insights). All NGS reads were trimmed, quality controlled, and aligned to a reference to confirm the DNA sequence.

### Primary CD4^+^ T cells

Human peripheral blood mononuclear cells (PBMCs) were isolated from buffy coats from anonymized healthy blood donors by Ficoll density centrifugation using Lymphoprep (Stemcell Technologies). Buffy coats were provided by the Blood Donation Service Zurich, Swiss Red Cross in Schlieren, Switzerland, and blood donors gave written informed consent for the use of buffy coats for research purposes. Enrichment of primary CD4^+^ T cells was performed using the EasySep™ Human CD4^+^ T Cell Enrichment Kit (Miltenyi Biotec) according to the manufacturer’s manual. Primary CD4^+^ T cells were kept in culture using RPMI-1640 medium supplemented with L-glutamine, 10% heat-inactivated FBS, 1% P/S and 50 U/mL IL-2 (Roche), at 37°C and 5% CO_2_.

Prior to Ad transduction, CD4^+^ T cells were activated with Dynabeads™ Human T-Activator CD3/CD28 (Gibco by Thermo Fisher Scientific) according to the manufacturer’s manual. After 24 h cells were separated from the beads over a magnetic column and several washing steps with pre-warmed medium without IL-2 supplementation. Cells were centrifuged at 150 × g for 10 min and counted.

### Adenovirus retargeting and transduction

Adenoviruses (Ads) containing either the CRISPR-HIV1, CRISPR-Con or iRFP reporter expression cassettes were preincubated with CD3-, CD28- and IL-2-retargeting adapters in a 32.5-fold total molar excess over adenoviral fiber knob for 1.5 h on ice in PBS (10.8-fold molar excess of each single adapter). Production and purification of adapters retargeting adapters was performed as previously described ^20^. 1×10^5^ activated primary CD4^+^ T cells were seeded in a 96-well plate and transduced with 2×10^4^ VP (virus particles)/cell of retargeting adapter-coated Ad-iRFP, Ad-CRISPR-HIV1 or Ad-CRISPR-Con. Cells were cultured in media without IL-2 supplementation.

### HIV-1 reporter virus preparation and infection

HIV-1_NL4-3-GFP Nef+_ virus stock was generated by transfecting HEK 293T cells of at least 90% confluency in a T-75 flask with the following components: pBR-NL4-3 Nef^+^ IRES eGFP vector (25 µg) and polyethylenimine (50 μg), and serum-free DMEM. Medium was replaced 16 h post-transfection. Supernatant-containing viral particles were harvested 48 h post-transfection, centrifuged at 600 × g for 10 min and the cell-free supernatant was sterile-filtered (0.22 µm) before freezing virus stock aliquots at -80°C. Virus stock was titrated by preparing a 1:5 dilution series of the virus stock in a 96-well plate and performing a TZM-bl assay with 1×10^4^ TZM-bl cells/well and measuring luciferase expression 72 h later. Ad transduced primary CD4^+^ T cells were infected with HIV-1_NL4-3-GFP Nef+_ virus with a MOI 0.05 and spinoculation of cells with the virus at 1200 × g for 2 h at 25°C. Afterwards, cells were resuspended in fresh media supplemented with 50 U/mL IL-2 and transferred back to the culture plate.

### Flow cytometry

Ad transduced and HIV-1_NL4-3-GFP Nef+_ infected primary CD4^+^ T cells were analyzed by flow cytometry 48 h post Ad transduction/24 h post HIV-1 infection. Cells were washed twice with PBS, fixed with 2% paraformaldehyde, and fluorophore expression was measured in a minimum of 10,000 events per sample with a BD LSR II Fortessa 4L (BD Biosciences). HIV-1- and/or Ad-positive were analyzed with FlowJo software v10.0.8.

### Statistical analysis

The data are presented as the mean and standard deviation (SD) of independent experiments. Statistical analysis was performed using Prism 9 software (Graph Pad, San Diego, CA). Ordinary one-way ANOVA tests were performed comparing the respective HIV-1-specific gRNA transfected samples to ‘No gRNA’ control to evaluate the statistical significance of the observed effects. Alternatively, paired, two-tailed t-tests were performed comparing samples where indicated. A result of P <0.033 was considered to be statistically significant. P <0.033 is indicated with *, P <0.002 with **, P <0.0002 with *** and P <0.0001 with ****.

## Results

### Selection of HIV-1 specific gRNAs

The first step towards the successful application of CRISPR/Cas9 against HIV-1 entails the selection of suitable target sequences in the viral genome. The selection of HIV-1-specific gRNAs for our study was based on previous studies focusing on CRISPR/Cas9 targeting and disruption of HIV-1 that were published until 2018 (Table 2 and Fig. 1a). We screened the literature comparing the tested HIV-1-specific gRNA target sites with regard to their efficacy to inhibit HIV-1 in the respective gRNA screening systems, and selected six gRNAs in total for our study (Table 2). Four of the selected gRNAs target the HIV-1 LTRs and two target the *gag* gene, and all showed very effective HIV-1 inhibition in one or more of the previous studies (Table 2). Furthermore, we aligned each selected gRNA sequence to the Los Alamos National Laboratory (LANL) HIV sequence database to determine the HIV-1 sequence conservation of the target sites, because HIV-1 sequence diversity is a major challenge for HIV-1 therapies in general and a CRISPR/Cas9-based HIV-1 therapy in particular. The gRNAs –LTR2, –TAR and –Gag exhibited the highest sequence conservation with ∼20-30% among all HIV-1 subtypes, and ∼58-68 % among subtype B. In comparison, gRNA–p24 exhibited sequence conservation two-times lower and gRNAs –LTR1 and –LTR3 three-times lower (Table 2).

**Figure 1.**
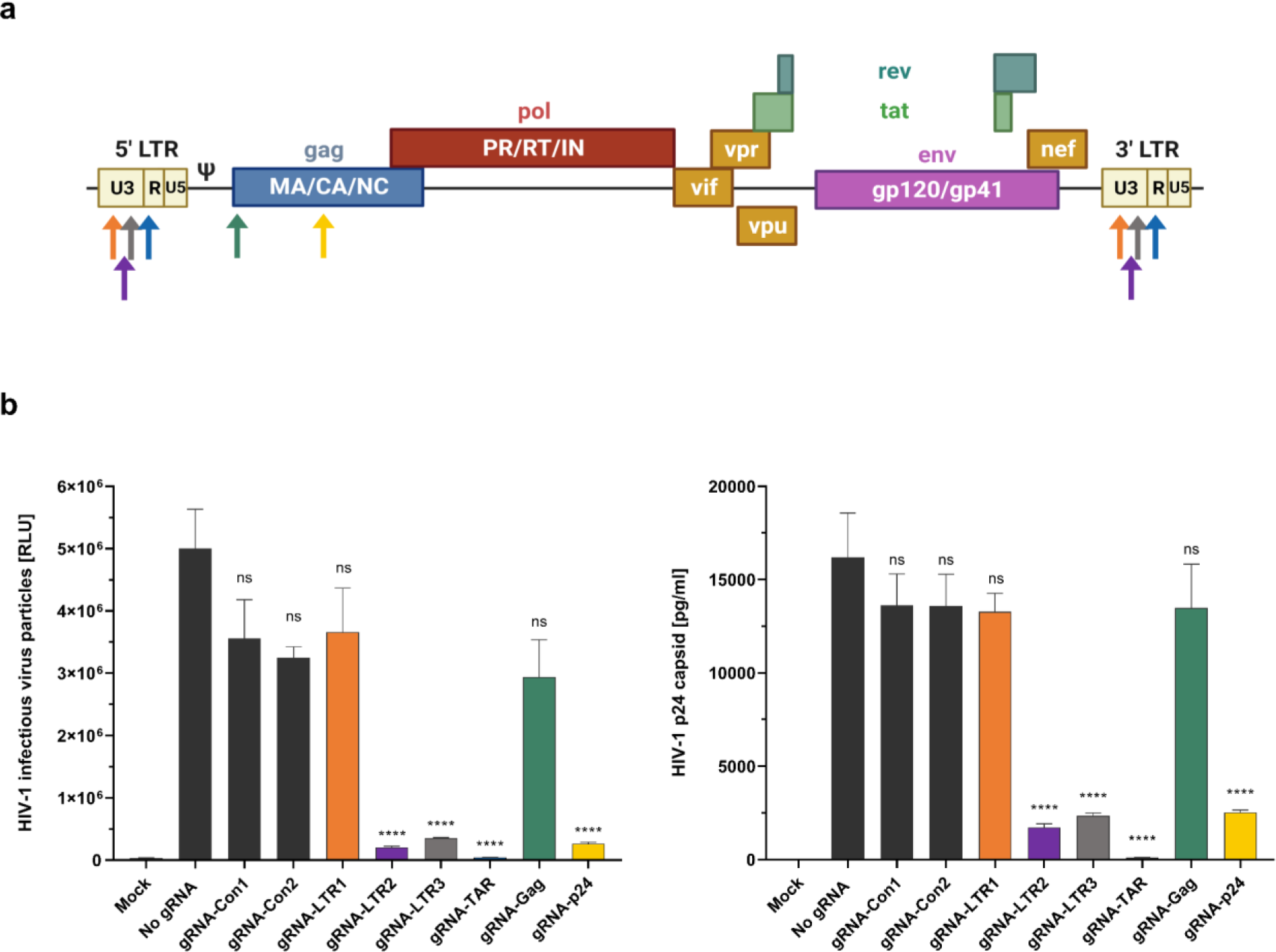
Inhibition of HIV-1 replication by CRISPR/Cas9. (a) Illustration of the HIV-1 genome and the HIV-1 targeting gRNAs. Six HIV-1 specific gRNAs were selected from different publications (Table 2). Four of these gRNAs target the HIV-1 LTRs and the other two gRNAs target *gag*. Figure created with BioRender.com (Figure publication license MO24KDZA3H). (b) 2×10^4^ HEK293T cells were seeded per well in a 96-well plate and 24 h later co-transfected with the pX458 plasmid containing the Cas9 nuclease gene and one of the selected HIV-1 gRNAs, as well as with an HIV-1 JR-CSF expression plasmid. pX458 without a gRNA (‘No gRNA’) was co-transfected with the HIV-1 JR-CSF expression plasmid as a control. The ‘Mock’ sample represents non-transfected cells. 48 h post transfection the cell supernatant was harvested and HIV-1 replication was measured by analyzing 50 µl cell supernatant for HIV-1 p24 capsid protein (pg/ml) by ELISA, and 50 µl for HIV-1 infectious virus particles (relative light units, RLU) by TZM-bl luciferase assay. Shown are means ±SD with n=3. *P<0.033, **P<0.002, ***P<0.0002 and ****P<0.0001 indicate statistical significance compared to ‘No gRNA’ control by ordinary one-way ANOVA.

**Table 2.**
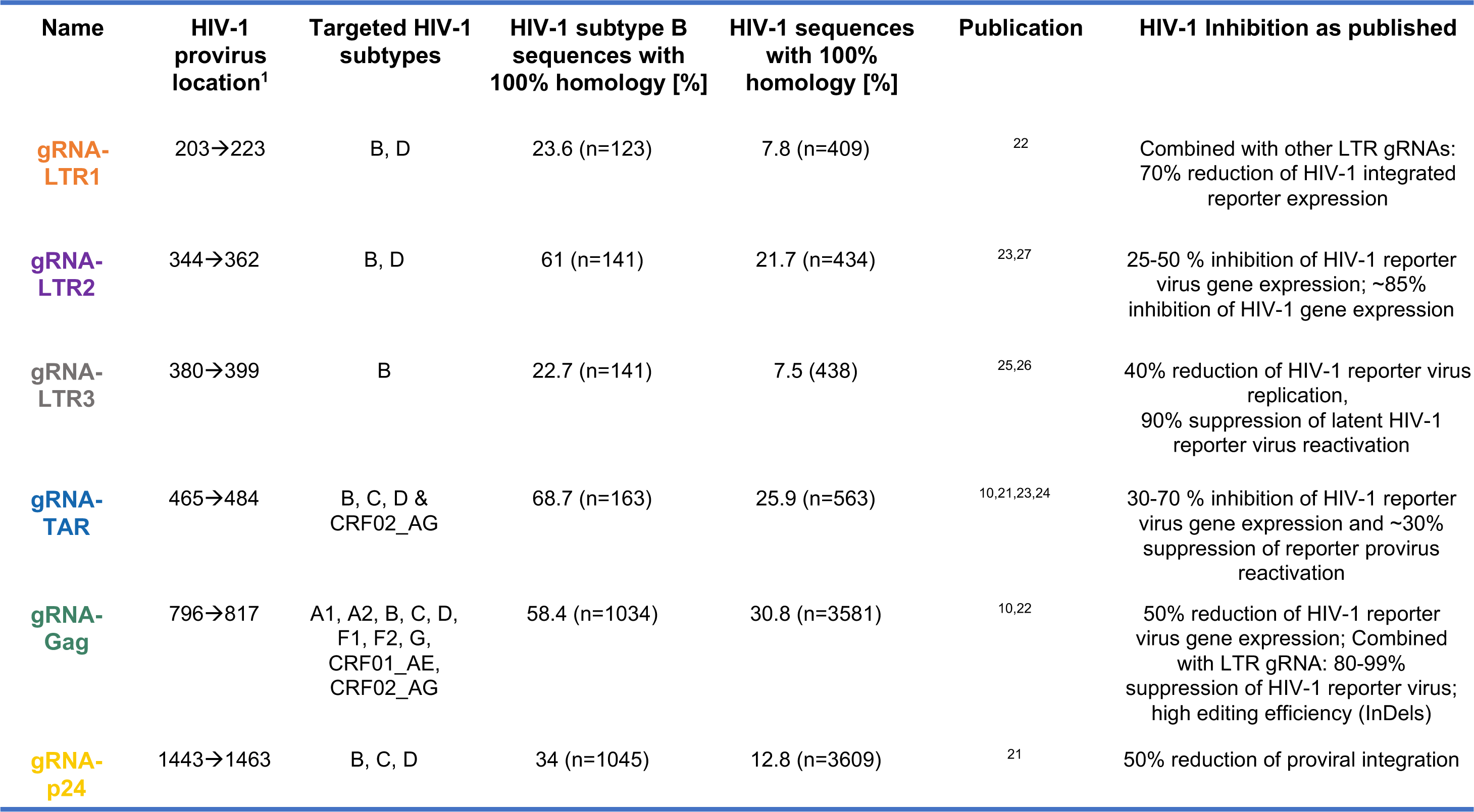
Selection of HIV-1 specific gRNA. HIV-1 specific gRNAs tested in this study were previously published and selected based on their efficacy to inhibit HIV-1 replication in the respective studies. gRNA sequences were aligned to all HIV-1 sequences in the Los Alamos National Laboratory (LANL) HIV sequence database (<u>HIV sequence database main page (lanl.gov)</u>) by using the QuickAlign tool. HIV-1 sequences with 100% homology to the gRNAs are shown as [%] of all aligned (n) HIV sequences or subtype B. Differences in aligned sequences (n) is due to differences in stored sequences in the database. CRF= circulating recombinant form. ^1^ based on HIV-1 HXB2 (Genbank asseccion number K03455.1)

### Validation of the inhibitory effect of the selected HIV-1 specific gRNAs

To test the selected HIV-1-specific gRNAs regarding their antiretroviral efficacy against HIV-1 we performed a co-transfection screen in human embryonic kidney (HEK293T) cells, which included plasmids encoding HIV-1 and SpCas9 in the absence (No gRNA, Con1 and Con2) or presence of HIV-1-specific gRNAs (Fig. 1b). The HIV-1 subtype B molecular clone JR-CSF was used as well as two random nucleotide (nt) gRNA controls, gRNA–Con1 and –Con2. Analysis of cell culture supernatants revealed that CRISPR/Cas9 combined with gRNAs – LTR2, –LTR3, –TAR or –p24 significantly reduced the HIV-1 virion production, whereas the gRNA controls and gRNA–LTR1 and –Gag showed no effect on HIV-1 particle production (Fig. 1b). The effective gRNAs, –LTR2, –LTR3, –TAR or –p24, reduced the amount of infectious HIV-1 in supernatants by more than 90% compared to the No gRNA control, and gRNA–TAR showed the most potent effect, reducing HIV-1 by >99% in both assays (Fig. 1b).

### Efficient inhibition of HIV-1 through the combination of HIV-1 specific gRNAs

Next, we assessed whether combination of HIV-1-specific gRNAs could have a synergistic effect and increase antiretroviral efficacy of the CRISPR/Cas9 system in the HEK 293T cell co-transfection screen.

Combination of the effective gRNAs –LTR2, –LTR3, –TAR and –p24 with each other in most cases further increased the reduction of HIV-1 virion production compared to the respective gRNAs alone. For instance, combination of gRNA–LTR2 with gRNA–LTR3, –TAR or –p24 revealed a significant decrease of HIV-1 virion production compared to gRNA–LTR2 alone indicating a synergistic effect of these gRNAs (Fig. 2). Combination of gRNA–LTR3 or –p24 with gRNA–LTR2 or –TAR also exhibited a significant reduction compared to gRNA–LTR3 or gRNA–p24 alone (Fig. 2).

**Figure 2.**
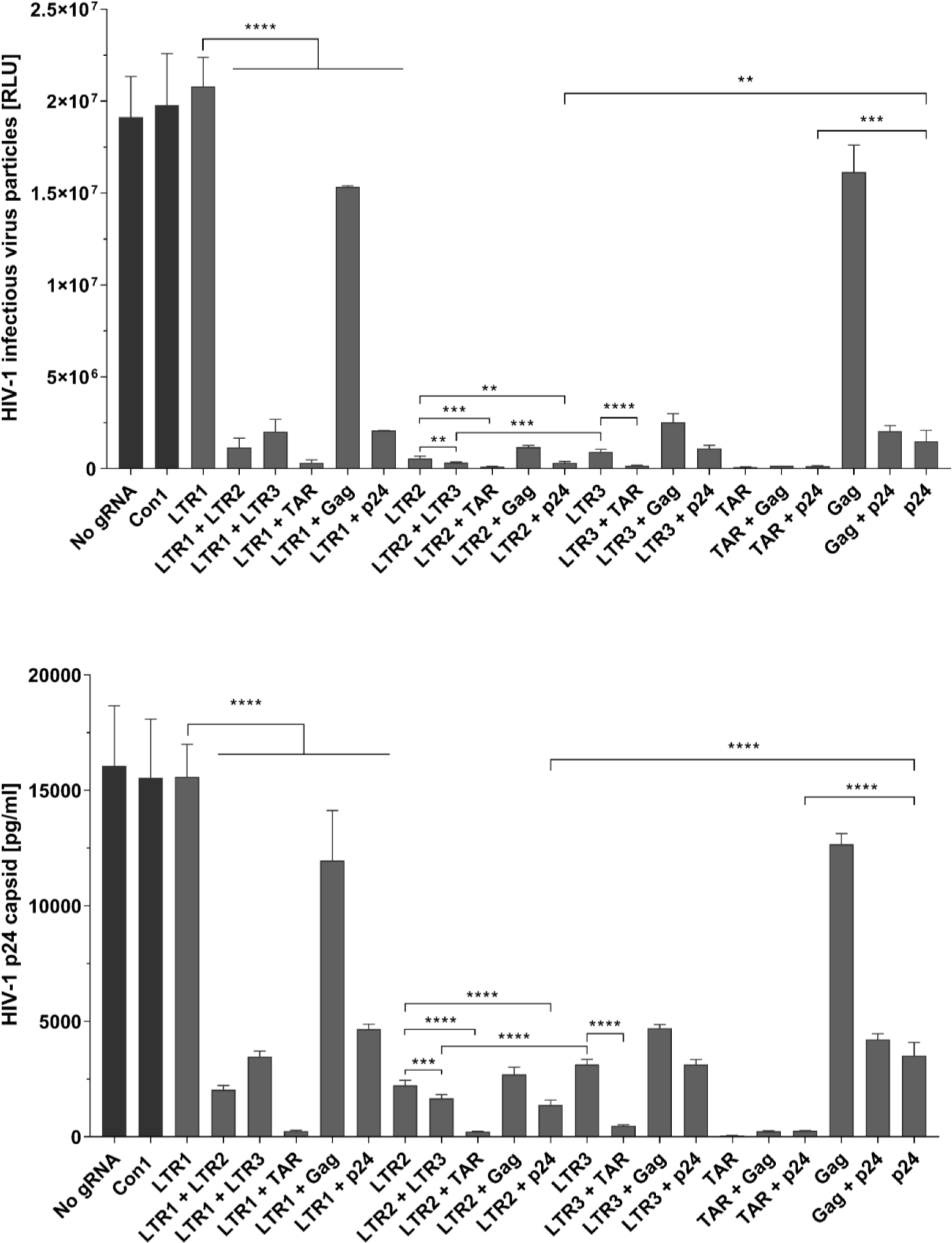
Robust inhibition of HIV-1 by gRNA–TAR editing in combination with less effective gRNAs. 2×10^4^ HEK293T cells were seeded per well in a 96-well plate and 24 h later co-transfected with two pX458 plasmids containing different HIV-1 gRNAs and the Cas9 gene, as well as with an HIV-1 JR-CSF expression plasmid in a 2.5:2.5:1 mass ratio. pX458 without a gRNA (‘No gRNA’) was co-transfected with the HIV-1 JR-CSF expression plasmid. 48 h post transfection the cell supernatant was harvested and HIV-1 replication was measured by analyzing 50 µl cell-free supernatant for HIV-1 p24 capsid protein (pg/ml) by ELISA, and 50 µl for HIV-1 infectious virus particles (relative light units, RLU) by TZM-bl luciferase assay. Shown are means ±SD with n=3. *P<0.033, **P<0.002, ***P<0.0002 and ****P<0.0001 indicate statistical significance of gRNA combination compared to the indicated single gRNA by ordinary one-way ANOVA.

The gRNA–TAR was uncovered as the most potent and robust gRNA, because its combination with any other gRNA showed a consistent reduction in HIV-1 virion production by >99%, also when combined with the ineffective gRNAs –LTR1 or –Gag. Vice versa, its effect could not further be improved by combination with the other selected gRNAs (Fig.2).

Also the other effective gRNAs –LTR2, –LTR3, or –p24, displayed a strong >85% reduction of HIV-1 virion production when combined with the ineffective gRNAs –LTR1 or –Gag, demonstrating the antiretroviral efficacy and capability of the effective gRNAs to compensate ineffective gRNAs.

**The combination of gRNA-TAR/p24 efficiently leads to large deletions in the HIV-1 DNA** A dual gRNA approach to combating HIV-1 is advantageous because, for example, it can induce more deleterious mutations and even larger deletions. We selected gRNA–p24 for combination with gRNA–TAR, because of its efficacy and because it targets the *gag* gene that encodes for HIV-1 structural proteins., The combination of gRNA–TAR and –p24 (Fig. 3a) induced a ∼1 kb deletion that was detectable by PCR and next generation sequencing (NGS) (Fig. 3b and c). Gel electrophoresis revealed two bands, the smaller one representing the “truncated (trunc-) LTR/Gag” amplicon with the deletion between the gRNA target sites. NGS results of the trunc-LTR/Gag amplicon confirmed the deletion (Fig. 3c). The full-length PCR product “5’LTR-Gag” was separately analyzed by NGS and deletions at the expected Cas9-induced break points were observed (Fig. 3b).

**Figure 3.**
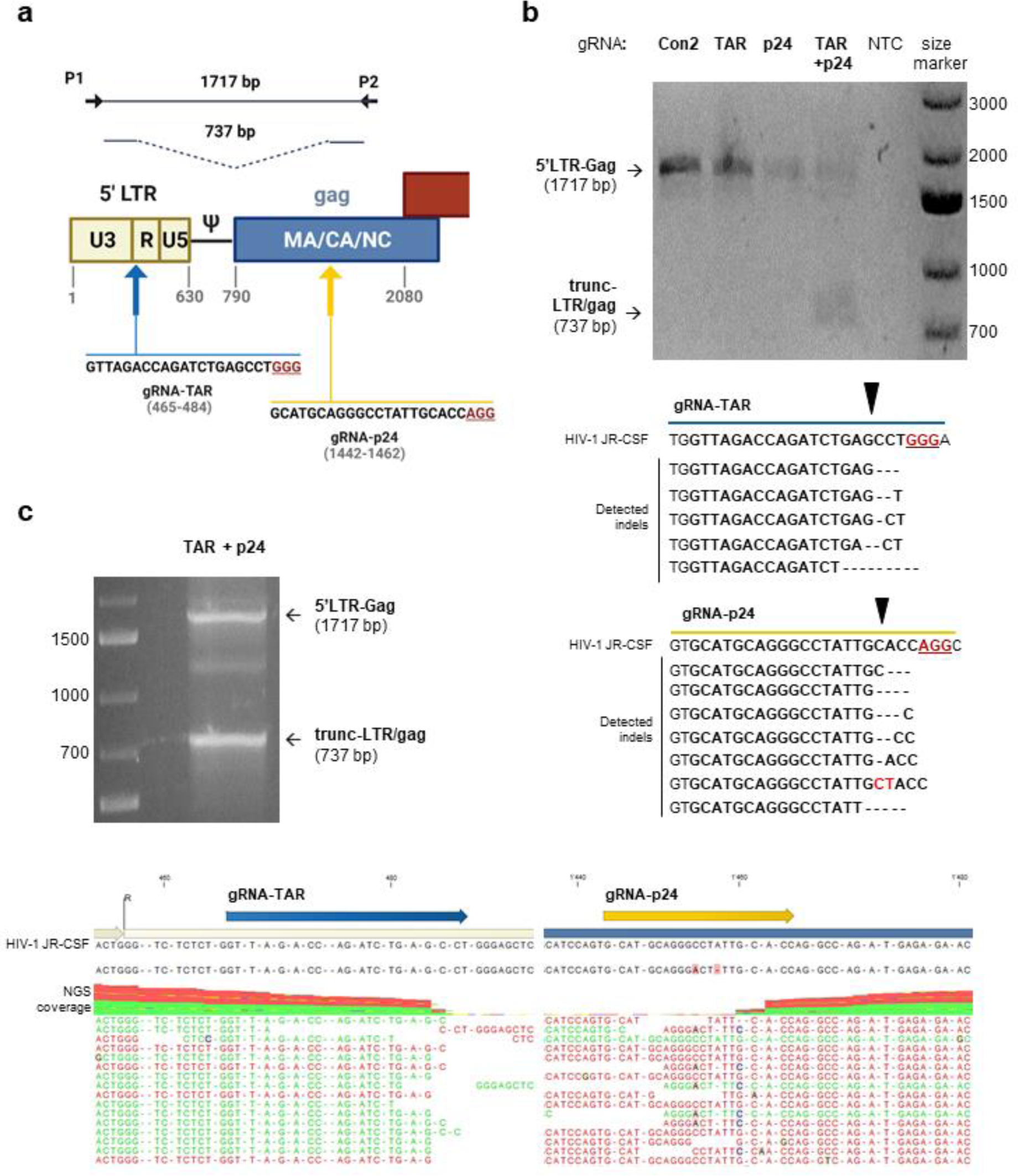
Editing of HIV-1 DNA by CRISPR/Cas9 and gRNAs TAR and p24. (a) Illustration of the 5’ region of the HIV-1 genome and the gRNA–TAR and –p24 target sites and sequences. The designed InDel PCR with primers P1 and P2 encloses both gRNA target sites. Predicted PCR amplicons are shown. Figure created with BioRender.com (Figure publication license MO24KDZA3H). (b) 1×10^5^ HEK 293T cells were seeded per well in a 12-well plate and 24 h later co-transfected with one, (Con2, TAR or p24), or two, (TAR and p24), pX458 plasmids, containing the Cas9 gene and the respective HIV-1 gRNAs. The HIV-1 JR-CSF expression plasmid was also co-transfected in a 1:5 or 1:2.5:2.5 mass ratio to the pX458 plasmids. 48 h post transfection cells were pelleted, washed with PBS, and a DNase treatment was performed before total genomic DNA was isolated. PCR with P1 and P2 was performed with the isolated DNA and the predicted amplicons, “5’LTR-Gag” or “trunc-LTR/gag”, and their sizes are shown as a gel electrophoresis analysis of the PCR reactions. Amplicons were purified and next-generation sequencing (NGS) performed. Analysis of NGS reads detected InDels at the gRNA target sites. Cas9 break point is indicated by a black arrow. (c) Gel electrophoresis analysis of the PCR reaction of the dual gRNA treated cells is shown. Bands were cut from gel, purified and separately subjected to NGS. Analysis of the NGS results of the truncated (trunc-LTR/gag) amplicon confirmed a 1 kb deletion between the gRNA target sites, showing a strong drop in NGS read coverage at the Cas9 break points.

### HIV-1 inhibition by gRNA-TAR/p24 delivered by CD3-CD28-IL2-retargeted adenovirus vector in primary CD4^+^ T cells

Next, we aimed to demonstrate the suppression of HIV replication in human primary CD4^+^ T cells (Fig. 4b). To achieve high transduction efficiency, we delivered the HIV-1 specific gRNAs through novel T cell specific CD3-CD28-IL2-retargeted Adenovirus (Ad) vector ^20,28^. Ad-CRISPR-HIV1 containing the SpCas9 and both selected HIV-1-specific gRNAs, –TAR and –p24, were tested in comparison to Ad-CRISPR-Con, which contained the two random nucleotides gRNAs –Con1 and –Con2 tested previously (Fig. 4a). Primary CD4^+^ T cells from uninfected donors were transduced with Ad-CRISPR-HIV1 or Ad-CRISPR-Con and afterwards infected with the HIV-1_NL4-3-GFP Nef+_ infectious reporter virus. By using an Ad-iRFP reporter virus and a HIV-1 GFP reporter virus, we were able to not only measure Ad transduction but also HIV-1 infection as well as co-infection of both via flow cytometry. The CD3-CD28-IL2-retargeted Ad-iRFP virus alone showed transduction of 62.3±23.3% of primary CD4^+^ T cells from three independent donors (Fig. 5b). The HIV-1 GFP reporter virus alone infected 11.8±9.2% of primary CD4^+^ T cells from three independent donors (Fig. 5c). Combination of both viruses revealed increased Ad transduction up to 75.6±5.9% as well as increase HIV-1 infection up to 20.2±5.8% (Fig. 4c and 5a). Most importantly, the majority of HIV-1-infected cells were also transduced with adenovirus, namely 80.4±7.7%, which demonstrated that the HIV-1-specific CRISPR/Cas9 system is expressed in HIV-1 target cells in our experimental setup (Fig. 5a).

**Figure 4.**
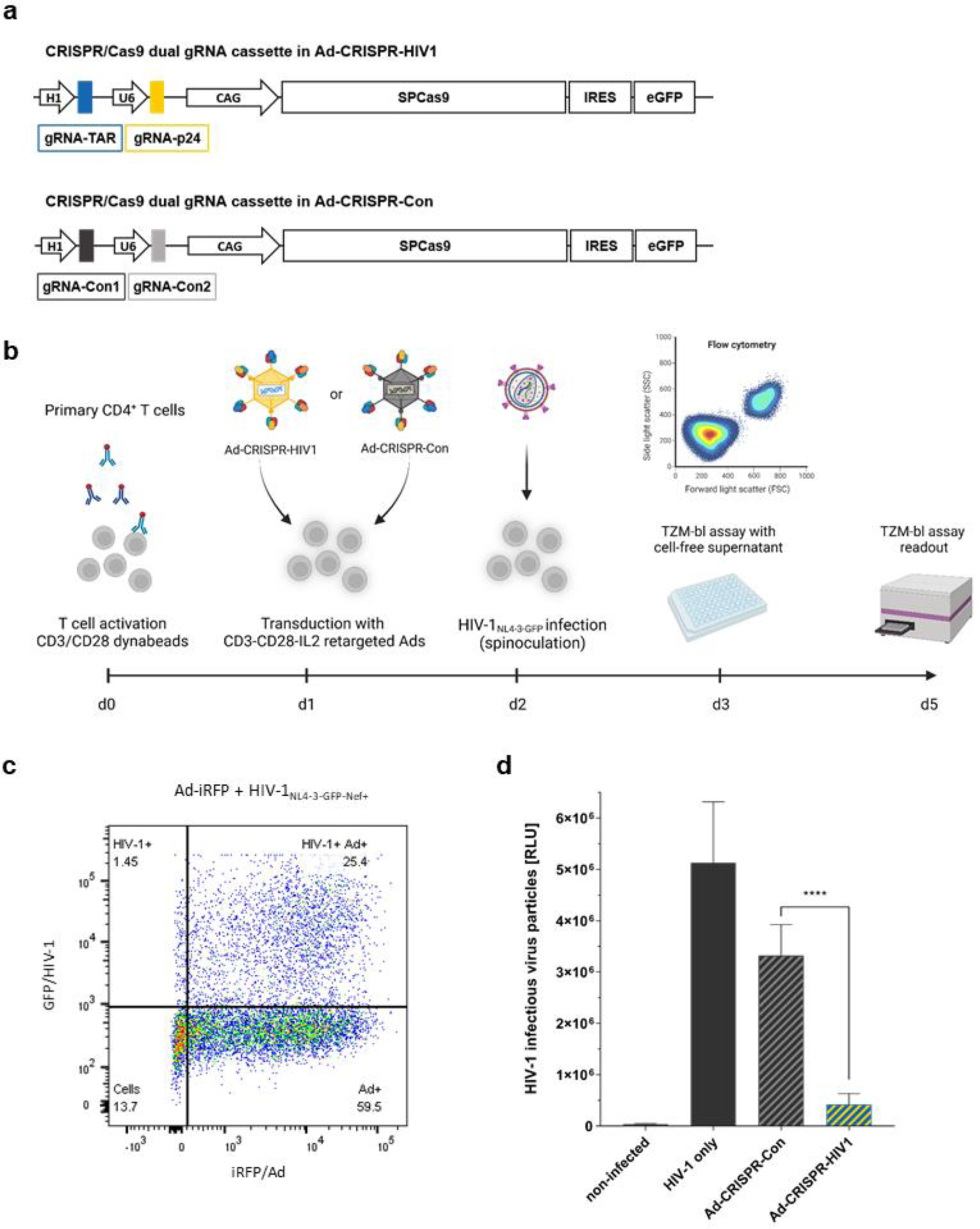
Retargeted Ad-CRISPR-HIV1 inhibits HIV-1 replication in primary CD4^+^ T cells. (a) CRISPR/Cas9 dual gRNA constructs cloned into Ad vectors are depicted. The Ad-CRISPR-HIV1 vector contains two expression cassettes for the HIV-1 specific gRNAs, –TAR and –p24. Downstream of it is the Cas9 and eGFP expression cassette. The Ad-CRISPR-Con control Ad is almost identical, but it contains the control gRNAs –Con1 and –Con2, which are random nucleotide sequences that are non-targeting. (b) The experimental workflow for Ad-CRISPR testing in primary CD4^+^ T cells is depicted. Figure created with BioRender.com (Figure publication license QP24J3WMYD). 10-15×10^6^ primary CD4^+^ T cells isolated from one donor were activated with CD3/CD28 Dynabeads for 24 h and after detachment from beads and washing with medium without IL-2, 1×10^5^ activated primary CD4^+^ T cells per well were seeded in a 96-well plate. Cells were transduced with 2×10^4^ VP (virus particles)/cell of retargeting adapter-coated Ad-iRFP (not depicted in scheme), Ad-CRISPR-HIV1 or Ad-CRISPR-Con. Ad coating was performed by preincubating Ads with CD3-, CD28- and IL-2-retargeting adapters ^20,28^ in a 32.5-fold total molar excess over adenovirus fiber knob for 1.5 h on ice before addition to cells. 24 h post Ad transduction cells were infected with HIV-1_NL4-3-GFP-Nef+_ virus (MOI 0.05) via spinoculation at 1200 × g for 2 h. After another 24 h the cell-free supernatant was used for TZM-bl assays and cells were fixed for flow cytometry analysis. (c) Ad transduction and HIV-1 infection efficiency with Ad-iRFP and HIV-1_NL4-3-GFP-Nef+_ reporter viruses in primary CD4^+^ T cells was measured by flow cytometry. Shown is an exemplary flow cytometry plot in primary CD4^+^ T cells from one donor exposing single- and double-positive cell populations. (d) HIV-1 replication was measured by analyzing 20 µl of cell-free supernatant for HIV-1 infectious virus particles (relative light units, RLU) by TZM-bl luciferase assays. The TZM-bl readout was performed 48 h after assay start. Shown are means ±SD of three independent experiments with primary CD4^+^ T cells from different donors. *P<0.033, **P<0.002, P<0.0002 and ****P<0.0001 indicate statistical significance between samples by paired, two-tailed t-tests.

**Figure 5.**
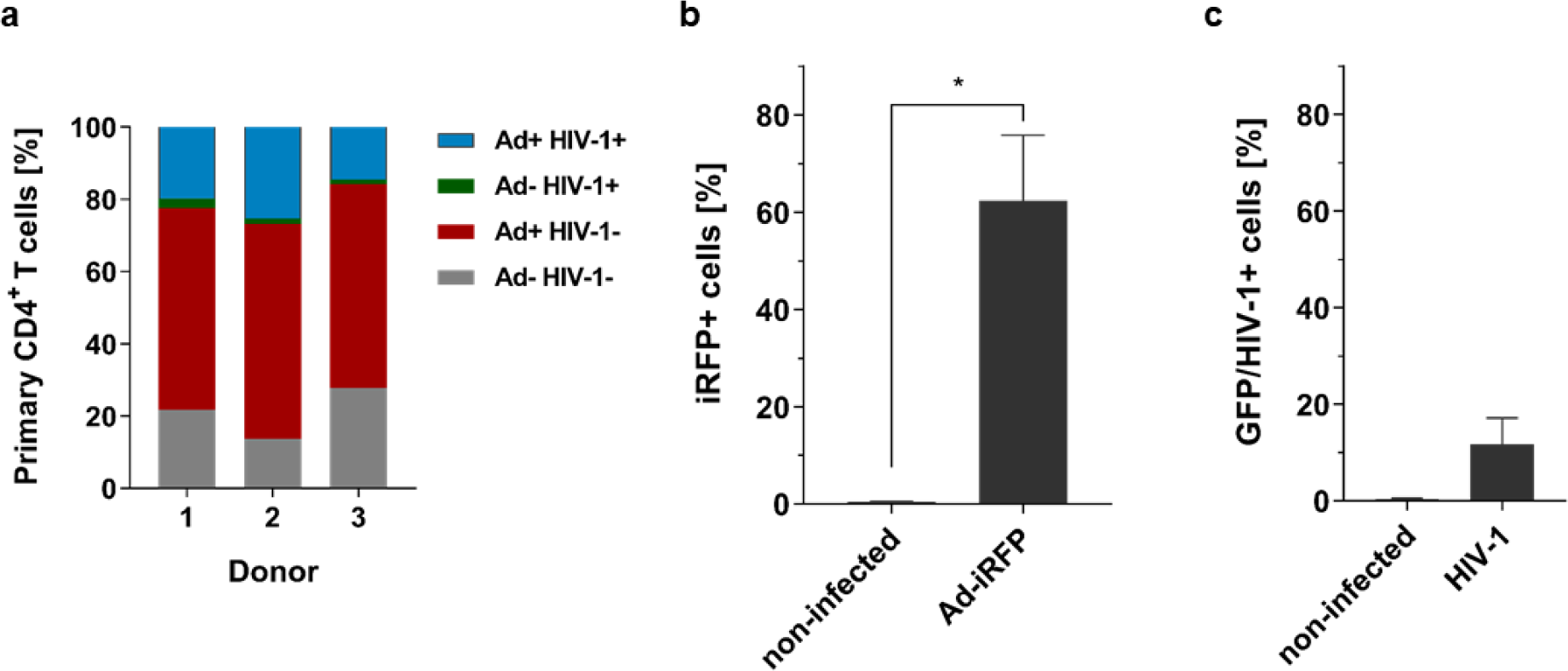
Ad transduction and HIV-1 infection efficiency in primary CD4+ cells. Flow cytometry analysis of CD3-CD28-IL2-retargeted Ad-iRFP reporter virus and HIV-1_NL4-3-GFP Nef+_ infected primary CD4+ T cells from different donors was performed on day 4. Shown are means ±SD of three independent experiments with primary CD4+ T cells from three different donors. *P<0.033, **P<0.002 and ***P<0.0002 indicate statistical significance between samples by paired, two-tailed t-test. (a) Composition of single-Ad or -HIV-1 positive and Ad-HIV-1-double positive primary CD4+ T cell populations. (b) Ad transduction efficiency shown with iRFP+ cells [%] only transduced cells and (c) HIV-1 infection shown with HIV-1_NL4-3-GFP Nef+_ only treated cells as GFP/HIV-1+ cells [%].

Finally, analysis of the cell supernatant confirmed that the HIV-1-targeting CRISPR/Cas9 system was successfully delivered by the CD3-CD28-IL2-retargeted Ad vector showing a 88.0±4.5% reduction of infectious HIV-1 virions compared to the Ad-CRISPR-Con control (Fig. 4d). Nonetheless, editing of HIV-1 DNA could not be confirmed by InDel PCR and NGS (data not shown), which together suggests that the system introduced here inhibits HIV-1 replication by rapid elimination of edited HIV-1 DNA before integration into the host cell genome.

## Discussion

HIV-1 provirus excision or inactivation by genome editing technologies such as ZFNs, TALENs and the CRISPR/Cas system is one of the major HIV-1 eradication strategies. Besides targeting of the latent provirus, the CRISPR/Cas system has also been investigated as a therapeutic approach to immunize cells against HIV-1 infection and/or suppress viral replication ^29^.

Here, we applied the *Staphylococcus pyogenes* (Sp) Cas9 with a dual HIV-1-specific gRNA approach that is delivered with T cell-targeted adenovirus vectors (Ads) to inactivate HIV-1. The most effective and robust LTR-targeting gRNA–TAR was multiplexed with the highly effective *gag* gene-targeting gRNA–p24, which when combined induced a ∼1 kb deletion in the HIV-1 DNA. Delivery of SpCas9 and both HIV-1-specific gRNAs by targeted Ads into primary CD4^+^ T cells was highly effective, transducing 62.3±23.3% of cells from three independent blood donors and inhibiting HIV-1 infection and replication by 88.0±4.5%.

Since the discovery of the CRISPR/Cas9 system it has been extensively studied regarding its application to inactivate HIV-1 ^29^. Identification of efficacious gRNA target sites that are conserved among divergent viral strains and multiplexing of gRNAs are key objectives for CRISPR/Cas therapy against HIV-1.

Numerous studies have shown that targeting of the HIV-1 LTRs with gRNAs, and specifically the U3 and R regions in the LTRs, leads to the strongest suppression of HIV-1 ^10,23,25,30-33^. Excision of the entire HIV-1 genome from the host cell genome with a single LTR-targeting gRNA was also shown to be possible due to the sequence identities of the LTRs ^11,25^. The gRNA–TAR tested in this work and previous studies targets the TAR sequence in the LTR R region, which is positioned at the neck of the stem loop region of TAR and is critical for Tat– TAR complex formation and initiation of HIV-1 transcription ^10,21,23,24,34^. Its targeting of a vulnerable regulatory region in the LTR explains its efficacy to inactivate HIV-1 that we and others observed, moreover it has been shown that as a single gRNA, gRNA–TAR is also capable of excising the whole provirus from the host genome ^23^. We detected cleavage and editing by gRNA-TAR at the expected Cas9 cleavage sites -3 nts upstream of the PAM, and the editing profile we observed was similar to a previous report showing that deletions occur up to 7 bp upstream of the gRNA–TAR PAM ^24,35,36^.

Multiplexing of gRNA–TAR has previously shown to improve HIV-1 disruption and excision ^10^. While an increase in HIV-1 suppression by gRNA–TAR in combination with other gRNAs was not apparent or rather not significantly improvable in our study, the combination with gRNA–p24 induced a ∼1 kb deletion between both gRNA target sites while showing the same level of efficacy. The gRNA–p24 targets the region that encodes the cyclophilin A-binding region of the p24 capsid protein ^37,38^. In an *in vitro* CRISPR digestion assay of a different study the gRNA–p24 exhibited the highest HIV-1 cleavage efficiency with 98.5%, while gRNA–TAR showed 50% cleavage efficiency ^21^. Two of the selected gRNAs, gRNA– LTR1 and –Gag showed no significant inhibition of HIV-1 in our co-transfection assay, whereas they showed an effect in previous studies ^10,22^. One explanation for the difference in our results is that we used SpCas9 to target these sites whereas C. Yin *et al.* applied the saCas9 to test both gRNAs ^22^. The effective combination of gRNA–TAR and –p24 is in line with studies that suggested combinations of an LTR-targeting gRNA with a gRNA that targets regions encoding for viral structural proteins, e.g. *gag* or *pol*, being more effective in eradicating HIV-1 genomes than LTR-targeting gRNAs alone ^26^. This approach showed promising results in a proof-of-concept *in vivo* study with *Staphylococcus aureus* (sa) Cas9 delivered by recombinant adeno-associated virus (rAAV) ^39^.

Here, we applied a novel CD3-retargeted adenovirus delivery method of the CRISPR/Cas system, which provides high T cell targeting and specificity ^20,28^. Delivery of SpCas9 and the HIV-1-specific gRNAs –TAR and –p24, as the T cell targeted Ad-CRISPR-HIV1, into primary CD4^+^ cells from three blood donors revealed strong inhibition of HIV-1 replication. While previous studies reported a reduction of virus production by more than three-fold with gRNA– TAR alone in primary CD4^+^ T cells, we observed an almost 9-fold or 88.0±4.5% reduction with gRNA–TAR and –p24 ^10^. Because the Cas9 nuclease and gRNAs –TAR and –p24 were already delivered and expressed in cells before challenge with HIV-1, the HIV-1 DNA was rapidly disrupted and probably eliminated before integration into the cell genome, which is why InDels could not be detected. Thus, the CRISPR/Cas9 system behaved as an HIV-1 defense system and is able to edit integrated as well as non-integrated target sequences as suggested previously ^10,11,21,25,40^. The next step will be to test it *ex vivo* to determine whether it is comparable to a previous lentivirus-delivered dual-LTR-gRNA CRISPR/Cas9 approach that showed 62-92% reduction of viral copy numbers in PBMCs and primary CD4^+^ cells of HIV-1 infected individuals ^11^.

A challenge for a CRISPR/Cas-based HIV-1 therapy is the virus sequence diversity that needs to be covered by gRNAs. The gRNA–TAR is relatively conserved among HIV-1 subtypes. Our sequence alignment to the LANL HIV database revealed complete matching of the gRNA– TAR sequence to 25.9% of all HIV sequences and 68.7% of all HIV-1 subtype B sequences present in the database ^41^. Among the non-B subtypes, gRNA–TAR also targets variants from subtypes C, D and the circulating recombinant (CRF) 02_AG. This is a minimal estimate of the HIV-1 diversity coverage by our tested gRNA since it only considers sequences included in the LANL HIV database ^42^. Furthermore, studies have shown that the 5’-proximal ∼10-12 nts of the gRNA are the “seed” region, which require 100% homology with the target sequence for DNA recognition, pairing and cleavage by the gRNA-Cas9 ribonucleoprotein complex, hence defining Cas9 specificity ^35,43^. Thus, HIV-1 variants that show mismatches outside of the seed region will be targeted by Cas9 but the more mismatches are present the lower the cleavage efficiency will be ^44^.

The importance of selection of highly conserved gRNA target sites and gRNA multiplexing is further stressed by the emergence of mutant viruses that are resistant to the CRISPR treatment. Treatment of cells with a single HIV-1-specific gRNA has shown to lead to the emergence of CRISPR-resistant mutant viruses, which exhibit resistance mutations at the gRNA target sequence ^21,24,30,45^. These escape mutations at the CRISPR/Cas target site partly arise from the error-prone reverse transcriptase and the host restriction factor APOBEC3G during HIV-1 replication; however, it could also be demonstrated that repeated Cas9 cleavage itself, and subsequent non-homologous end joining (NHEJ) of the Cas9-induced double-strand breaks (DSB) at the cleavage site, can cause escape mutations ^24,30,45^. Targeting highly conserved sequences and/or protein-coding sequences leads to a more sustained antiviral effect and longer delay in NHEJ-induced resistance mutations as shown previously for gRNA–TAR ^24,30,46^. Moreover, multiplexing of conserved LTR- and protein-coding-targeting gRNAs eliminates the emergence of CRISPR therapy escape ^46^.

Besides high gRNA target sequence conservation within the HIV-1 quasispecies, the ideal HIV-1 targeting gRNAs have no homology with the host genome to minimize CRISPR/Cas off-target cleavage, which is a concern with all genome editing technologies. As a retrovirus, HIV-1 shares similarity with human endogenous retroviruses (HERVs), which make up about 8% of the human genome, but only 5.6% exhibit even a low sequence similarity to the HIV-1 5’LTR sequence ^47-49^. A sophisticated gRNA design with bioinformatics tools is therefore able to avoid sequence homology to the human genome and prevent potential off-target effects ^49^. Thus far, numerous studies reported no severe off-target editing, cyto- or genotoxicity when using highly conserved gRNA target sequences ^11,25^.

The last challenge for a CRISPR/Cas editing or any gene therapy is the safe and efficient *in vivo* delivery. The most common *in vivo* delivery approach are viral vectors such as lentiviruses, adenoviruses (Ad) and especially adeno-associated adenoviruses (AAV). Although AAVs have been predominantly applied in CRISPR/Cas clinical trials due to their perceived lower immungenicity, adenoviral vectors might be more advantageous for CRISPR/Cas9 therapies.

Adenovirus vectors offer a larger transgene capacity especially of the helper-dependent (“gutless”) type ^50^, tissue specific and efficient delivery as well as capsid shielding technologies to evade immune recognition ^51,52^. In addition, with the new retargeting technology for CD3, CD28 and IL-2 receptors that we applied here, a more specific and efficient delivery of the CRISPR/Cas9 therapy into T cells was achieved ^20,28^.

In summary, our targeted Ad-CRISPR-HIV1 approach demonstrated a high efficacy to suppress HIV-1 replication in primary CD4^+^ T cells with a high level of HIV-1 and T cell specificity. Further steps would be to test its potency to target and inactivate latent HIV-1 provirus.

## Acknowledgements

We would like to thank Feng Zhang for pSpCas9(BB)-2A-GFP (PX458) (Addgene plasmid #48138). The following reagent was obtained through the NIH AIDS Reagent Program, Division of AIDS, NIAID, NIH: pYK-JRCSF (I.S.Y. Chen and Y. Koyanagi), pBR-NL4-3 Nef+ IRES eGFP (F. Kirchhoff) and TZM-bl cells (J.C. Kappes and X. Wu).

## Author contributions

SK, PCF, AP, and KJM designed the study. SK, PCF, and KJM conceptualized the study. SK performed experiments. SK analyzed the data. SK and KJM wrote the manuscript. All authors have read, edited and agreed to the published version of the manuscript.

## References

1 TW Chun, D Finzi, J Margolick, K Chadwick, D Schwartz & RF Siliciano. In vivo fate of HIV-1-infected T cells: quantitative analysis of the transition to stable latency. Nat Med 1, 1284–1290, (1995).

2 DD Ho, AU Neumann, AS Perelson, W Chen, JM Leonard & M Markowitz. Rapid turnover of plasma virions and CD4 lymphocytes in HIV-1 infection. Nature 373, 123–126, (1995).

3 TW Chun, RT Davey, Jr., M Ostrowski, J Shawn Justement, D Engel, JI Mullins & AS Fauci. Relationship between pre-existing viral reservoirs and the re-emergence of plasma viremia after discontinuation of highly active anti-retroviral therapy. Nat Med 6, 757–761, (2000).

4 M Wayengera. Proviral HIV-genome-wide and pol-gene specific zinc finger nucleases: usability for targeted HIV gene therapy. Theor Biol Med Model 8, 26, (2011).

5 X Qu, P Wang, D Ding, L Li, H Wang, L Ma, X Zhou, S Liu, S Lin, X Wang, G Zhang, S Liu, L Liu, J Wang, F Zhang, D Lu & H Zhu. Zinc-finger-nucleases mediate specific and efficient excision of HIV-1 proviral DNA from infected and latently infected human T cells. Nucleic Acids Res 41, 7771–7782, (2013).

6 H Ebina, Y Kanemura, N Misawa, T Sakuma, T Kobayashi, T Yamamoto & Y Koyanagi. A High Excision Potential of TALENs for Integrated DNA of HIV-Based Lentiviral Vector. PLOS ONE 10, e0120047, (2015).

7 M Romito, A Juillerat, YL Kok, M Hildenbeutel, M Rhiel, G Andrieux, J Geiger, C Rudolph, C Mussolino, A Duclert, KJ Metzner, P Duchateau, T Cathomen & TI Cornu. Preclinical Evaluation of a Novel TALEN Targeting CCR5 Confirms Efficacy and Safety in Conferring Resistance to HIV-1 Infection. Biotechnology Journal 16, 2000023, (2021).

8 K Khalili, R Kaminski, J Gordon, L Cosentino & W Hu. Genome editing strategies: potential tools for eradicating HIV-1/AIDS. J Neurovirol 21, 310–321, (2015).

9 J Herskovitz, M Hasan, M Patel, BD Kevadiya & HE Gendelman. Pathways Toward a Functional HIV-1 Cure: Balancing Promise and Perils of CRISPR Therapy. 27 (Springer US, 2022).

10 H-K Liao, Y Gu, A Diaz, J Marlett, Y Takahashi, M Li, K Suzuki, R Xu, T Hishida, C-J Chang, CR Esteban, J Young & JCI Belmonte. Use of the CRISPR/Cas9 system as an intracellular defense against HIV-1 infection in human cells. Nat Commun 6, 6413, (2015).

11 R Kaminski, Y Chen, T Fischer, E Tedaldi, A Napoli, Y Zhang, J Karn, W Hu & K Khalili. Elimination of HIV-1 Genomes from Human T-lymphoid Cells by CRISPR/Cas9 Gene Editing. Scientific Reports 6, 22555, (2016).

12 FA Ran, PD Hsu, J Wright, V Agarwala, DA Scott & F Zhang. Genome engineering using the CRISPR-Cas9 system. Nature protocols 8, 2281–2308, (2013).

13 S Konermann, MD Brigham, AE Trevino, J Joung, OO Abudayyeh, C Barcena, PD Hsu, N Habib, JS Gootenberg, H Nishimasu, O Nureki & F Zhang. Genome-scale transcriptional activation by an engineered CRISPR-Cas9 complex. Nature 517, 583–588, (2015).

14 Y Koyanagi, S Miles, RT Mitsuyasu, JE Merrill, HV Vinters & IS Chen. Dual infection of the central nervous system by AIDS viruses with distinct cellular tropisms. Science 236, 819–822, (1987).

15 M Schindler, J Münch & F Kirchhoff. Human immunodeficiency virus type 1 inhibits DNA damage-triggered apoptosis by a Nef-independent mechanism. J Virol 79, 5489–5498, (2005).

16 M Schmid, P Ernst, A Honegger, M Suomalainen, M Zimmermann, L Braun, S Stauffer, C Thom, B Dreier, M Eibauer, A Kipar, V Vogel, UF Greber, O Medalia & A Plückthun. Adenoviral vector with shield and adapter increases tumor specificity and escapes liver and immune control. Nat Commun 9, 450, (2018).

17 RB DuBridge, P Tang, HC Hsia, PM Leong, JH Miller & MP Calos. Analysis of mutation in human cells by using an Epstein-Barr virus shuttle system. Mol Cell Biol 7, 379–387, (1987).

18 X Wei, JM Decker, H Liu, Z Zhang, RB Arani, JM Kilby, MS Saag, X Wu, GM Shaw & JC Kappes. Emergence of Resistant Human Immunodeficiency Virus Type 1 in Patients Receiving Fusion Inhibitor (T-20) Monotherapy. Antimicrobial Agents and Chemotherapy 46, 1896, (2002).

19 A Trkola, AB Pomales, H Yuan, B Korber, PJ Maddon, GP Allaway, H Katinger, CF Barbas, 3rd, DR Burton, DD Ho & et al. Cross-clade neutralization of primary isolates of human immunodeficiency virus type 1 by human monoclonal antibodies and tetrameric CD4-IgG. J Virol 69, 6609-6617, (1995).

20 PC Freitag, M Kaulfuss, L Fluhler, J Mietz, F Weiss, D Brucher, J Kolibius, KP Hartmann, SN Smith, C Munz, O Chijioke & A Pluckthun. Targeted adenovirus-mediated transduction of human T cells in vitro and in vivo. Mol Ther Methods Clin Dev 29, 120–132, (2023).

21 S Ueda, H Ebina, Y Kanemura, N Misawa & Y Koyanagi. Anti-HIV-1 potency of the CRISPR/Cas9 system insufficient to fully inhibit viral replication. Microbiology and Immunology 60, 483–496, (2016).

22 C Yin, T Zhang, X Qu, Y Zhang, R Putatunda, X Xiao, F Li, W Xiao, H Zhao, S Dai, X Qin, X Mo, W-B Young, K Khalili & W Hu. In Vivo Excision of HIV-1 Provirus by saCas9 and Multiplex Single-Guide RNAs in Animal Models. Mol Ther 25, 1168–1186, (2017).

23 H Ebina, N Misawa, Y Kanemura & Y Koyanagi. Harnessing the CRISPR/Cas9 system to disrupt latent HIV-1 provirus. Scientific Reports 3, 2510, (2013).

24 KE Yoder & R Bundschuh. Host Double Strand Break Repair Generates HIV-1 Strains Resistant to CRISPR/Cas9. Scientific Reports 6, 29530, (2016).

25 W Hu, R Kaminski, F Yang, Y Zhang, L Cosentino, F Li, B Luo, D Alvarez-Carbonell, Y Garcia-Mesa, J Karn, X Mo & K Khalili. RNA-directed gene editing specifically eradicates latent and prevents new HIV-1 infection. Proc Natl Acad Sci 111, 11461–11466, (2014).

26 C Yin, T Zhang, F Li, F Yang, R Putatunda, WB Young, K Khalili, W Hu & Y Zhang. Functional screening of guide RNAs targeting the regulatory and structural HIV-1 viral genome for a cure of AIDS. AIDS 30, 1163–1174, (2016).

27 N Zhao, G Wang, AT Das & B Berkhout. Combinatorial CRISPR-Cas9 and RNA Interference Attack on HIV-1 DNA and RNA Can Lead to Cross-Resistance. Antimicrobial Agents and Chemotherapy 61, (2017).

28 S Klinnert, CD Schenkel, PC Freitag, HF Gunthard, A Pluckthun & KJ Metzner. Targeted shock-and-kill HIV-1 gene therapy approach combining CRISPR activation, suicide gene tBid and retargeted adenovirus delivery. Gene Ther, (2023).

29 J Herskovitz, M Hasan, M Patel, BD Kevadiya & HE Gendelman. Pathways Toward a Functional HIV-1 Cure: Balancing Promise and Perils of CRISPR Therapy. Methods Mol Biol 2407, 429–445, (2022).

30 G Wang, N Zhao, B Berkhout & AT Das. CRISPR-Cas9 Can Inhibit HIV-1 Replication but NHEJ Repair Facilitates Virus Escape. Mol Ther 24, 522–526, (2016).

31 Q Wang, S Liu, Z Liu, Z Ke, C Li, X Yu, S Chen & D Guo. Genome scale screening identification of SaCas9/gRNAs for targeting HIV-1 provirus and suppression of HIV-1 infection. Virus Research 250, 21–30, (2018).

32 Z Gao, M Fan, AT Das, E Herrera-Carrillo & B Berkhout. Extinction of all infectious HIV in cell culture by the CRISPR-Cas12a system with only a single crRNA. Nucleic Acids Res, (2020).

33 R Kaminski, Y Chen, J Salkind, R Bella, W-b Young, P Ferrante, J Karn, T Malcolm, W Hu & K Khalili. Negative Feedback Regulation of HIV-1 by Gene Editing Strategy. Scientific Reports 6, 31527, (2016).

34 S Richter, H Cao & TM Rana. Specific HIV-1 TAR RNA loop sequence and functional groups are required for human cyclin T1-Tat-TAR ternary complex formation. Biochemistry 41, 6391–6397, (2002).

35 M Jinek, K Chylinski, I Fonfara, M Hauer, JA Doudna & E Charpentier. A programmable dual-RNA-guided DNA endonuclease in adaptive bacterial immunity. Science 337, 816–821, (2012).

36 P Mali, L Yang, KM Esvelt, J Aach, M Guell, JE DiCarlo, JE Norville & GM Church. RNA-guided human genome engineering via Cas9. Science 339, 823–826, (2013).

37 T Hatziioannou, D Perez-Caballero, S Cowan & PD Bieniasz. Cyclophilin interactions with incoming human immunodeficiency virus type 1 capsids with opposing effects on infectivity in human cells. J Virol 79, 176–183, (2005).

38 GJ Towers. The control of viral infection by tripartite motif proteins and cyclophilin A. Retrovirology 4, 40, (2007).

39 R Kaminski, R Bella, C Yin, J Otte, P Ferrante, HE Gendelman, H Li, R Booze, J Gordon, W Hu & K Khalili. Excision of HIV-1 DNA by gene editing: a proof-of-concept in vivo study. Gene Ther 23, 690–695, (2016).

40 L Yin, S Hu, S Mei, H Sun, F Xu, J Li, W Zhu, X Liu, F Zhao, D Zhang, S Cen, C Liang & F Guo. CRISPR/Cas9 Inhibits Multiple Steps of HIV-1 Infection. Hum Gene Ther 29, 1264–1276, (2018).

41 LANL. Los Alamos HIV Sequence Database, <http://www.hiv.lanl.gov/> (

42 C Kuiken, B Korber & RW Shafer. HIV sequence databases. AIDS Rev 5, 52–61, (2003).

43 W Jiang, D Bikard, D Cox, F Zhang & LA Marraffini. RNA-guided editing of bacterial genomes using CRISPR-Cas systems. Nat Biotechnol 31, 233–239, (2013).

44 PD Hsu, DA Scott, JA Weinstein, FA Ran, S Konermann, V Agarwala, Y Li, EJ Fine, X Wu, O Shalem, TJ Cradick, LA Marraffini, G Bao & F Zhang. DNA targeting specificity of RNA-guided Cas9 nucleases. Nat Biotechnol 31, 827–832, (2013).

45 Z Wang, Q Pan, P Gendron, W Zhu, F Guo, S Cen, Mark A Wainberg & C Liang. CRISPR/Cas9-Derived Mutations Both Inhibit HIV-1 Replication and Accelerate Viral Escape. Cell Rep 15, 481–489, (2016).

46 G Wang, N Zhao, B Berkhout & AT Das. A Combinatorial CRISPR-Cas9 Attack on HIV-1 DNA Extinguishes All Infectious Provirus in Infected T Cell Cultures. Cell Rep 17, 2819–2826, (2016).

47 DJ Griffiths. Endogenous retroviruses in the human genome sequence. Genome Biology 2, reviews1017.1011, (2001).

48 AC van der Kuyl. HIV infection and HERV expression: a review. Retrovirology 9, 6, (2012).

49 RW Link, MR Nonnemacher, B Wigdahl & W Dampier. Prediction of Human Immunodeficiency Virus Type 1 Subtype-Specific Off-Target Effects Arising from CRISPR-Cas9 Gene Editing Therapy. The CRISPR Journal 1, 294–302, (2018).

50 D Brucher, N Kirchhammer, SN Smith, J Schumacher, N Schumacher, J Kolibius, PC Freitag, M Schmid, F Weiss, C Keller, M Grove, UF Greber, A Zippelius & A Pluckthun. iMATCH: an integrated modular assembly system for therapeutic combination high-capacity adenovirus gene therapy. Mol Ther Methods Clin Dev 20, 572–586, (2021).

51 P Boucher, X Cui & DT Curiel. Adenoviral vectors for in vivo delivery of CRISPR-Cas gene editors. Journal of Controlled Release 327, 788–800, (2020).

52 M Schmid, P Ernst, A Honegger, M Suomalainen, M Zimmermann, L Braun, S Stauffer, C Thom, B Dreier, M Eibauer, A Kipar, V Vogel, UF Greber, O Medalia & A Pluckthun. Adenoviral vector with shield and adapter increases tumor specificity and escapes liver and immune control. Nat Commun 9, 450, (2018).

